# Terrain ruggedness and human activities influence the distribution of Grauer’s gorillas (*Gorilla beringei graueri*) and eastern chimpanzees (*Pan troglodytes schweinfurthii*) in the Tayna Nature Reserve, Democratic Republic of Congo

**DOI:** 10.1101/2025.07.28.667236

**Authors:** Alice Michel, Jackson Kabuyaya Mbeke, Benezeth Kambale Visando, Sonya Kahlenberg, Katie Fawcett, Damien Caillaud

## Abstract

The Albertine Rift Hotspot in east-central Africa is one of the most biodiverse regions in the world. It hosts a number of conservation-priority species, including Critically Endangered Grauer’s gorillas and Endangered eastern chimpanzees. Yet, prolonged insecurity in the region has made wildlife monitoring and conservation challenging. This is particularly true of the northeastern limits of the range of Grauer’s gorillas, where wide stretches of unprotected forest could harbor behaviorally and genetically unique peripheral populations of great apes and other species. Here, we developed a rapid wildlife survey method to map population distributions and monitor the impact of various human activities in the Tayna Nature Reserve in eastern Democratic Republic of Congo. Despite a lack of surveys since the early 2000s, we find that Grauer’s gorillas and eastern chimpanzees, as well as a number of smaller-bodied species, have persisted in Tayna. Observations of gorilla and chimpanzee signs were highly heterogeneous. They were less dense near human settlements and artisanal mines and peaked in areas with rugged terrain. The association with rugged terrain may be a result of historical anthropogenic pressure, behavioral avoidance of human activities, and/or ecological preference. Maintaining connectivity between patches of suitable great ape habitat will be critical for long-term great ape conservation in the Albertine Rift region. Our findings also demonstrate the utility of priority species monitoring driven by local communities for conservation initiatives.

## 1. Introduction

Habitat loss, overhunting, and climate change are the predominant threats to terrestrial wildlife globally (Hogue and Breon, 2022). Yet wildlife populations vary in their response to human activities, depending on the type of activity, local context, and species. For example, whether hunting is sustainable depends on local ecological processes and prey behavioral responses (Ripple et al., 2016). In some cases, wildlife persists in or recolonizes areas with high hunting pressure (*e.g.*, African forest elephant, *Loxodontis cyclotis*, in a logging concession in Gabon; Van Vliet and Nasi, 2008; cougar, *Puma concolor,* in hunting areas in the USA; Robinson et al., 2008; African wild dog, *Lycaon pictus* in the buffer zone around Hwange National Park, Zimbabwe; van der Meer et al., 2014), while in other cases, wildlife avoids them (*e.g.,* African forest elephant in high human use areas within Dzanga-Sangha Protected Area Complex, Central African Republic; Blom et al., 2005; black bear, *Ursus americanus*, and elk, *Cervus canadensis*, in hunting areas in the USA; Stillfried et al., 2015; Spitz et al., 2019). Other activities that do not directly target wildlife may still cause avoidance, as for leopards (*Panthera pardus*) in pastoral lands in Kenya (Van Cleave et al., 2018) and chimpanzees (*Pan troglodytes*) in logging concessions in northern Congo (Morgan et al., 2018). Populations may even exhibit selective preference for human-dominated habitats: leopards in the Kalahari are more active near farms than protected areas (Van der Weyde et al., 2018) and chimpanzees raid crops in human-dominated landscapes (Hockings and McLennan, 2012; Krief et al., 2014). Finally, some species are ecological generalists (Ramiadantsoa et al., 2018) or specialists that thrive in secondary habitats created, for example, by selective logging, including wildlife targeted by hunting like western lowland gorillas (*Gorilla gorilla gorilla*; Morgan et al., 2018). Identifying how specific threats and habitat preferences interact at the population level is important to the design of conservation plans (Martin, 1998).

Tropical forests, the most biodiverse terrestrial biome, are facing escalating habitat degradation, forest fragmentation, and pressure from bushmeat hunting (Abernethy et al., 2013; Aleman et al., 2018; Taubert et al., 2018). The Albertine Rift in tropical east-central Africa includes 40% of the continent’s mammalian taxa in less than one percent of its total land area (Mittermeier et al., 2004). Though 60% of the region is under some degree of protection, it faces numerous conservation challenges, including armed conflicts, deforestation for charcoal, agriculture, mining, and oil extraction, and climate change (Plumptre et al., 2017). These processes interrelate, amplifying pressure on wildlife; for example, mining and logging increase hunting pressure in remote forests to feed workers, and hunters use new access roads that fragment forests to supply growing urban demand for bushmeat (Edwards et al., 2014; Laurance et al., 2006; Spira et al., 2019). As a result, keystone vertebrate species in the Albertine Rift, including forest elephants (*Loxodonta cyclotis*), forest buffalo (*Syncerus cafer nanus*), eastern chimpanzees (*Pan troglodytes schweinfurthii*), and gorillas (*Gorilla beringei*), have experienced population declines and local extirpations, with long-term consequences on forest ecosystem composition and structure (Mittermeier et al., 2004; Poulsen et al., 2018). In particular, the critically endangered Grauer’s gorilla (*Gorilla beringei graueri*), endemic to eastern Democratic Republic of Congo (DRC), declined from ∼17,000 individuals in the mid-1990s (Hall et al., 1998) to between 4,500 to 9,500 in 2019 (Plumptre et al., 2021, 2016). In addition to overall loss of numbers, the population has become fragmented (Plumptre et al., 2021, 2016). Sympatric endangered eastern chimpanzees have experienced a similar situation (Plumptre et al., 2021, 2016).

However, only a fraction of the range of Grauer’s gorillas has been surveyed in the past 20 years (Plumptre et al., 2021, 2016). In particular, there is a lack of data from the northeastern part of their range due to regional insecurity and logistical challenges (Hart and Hall, 1996; Plumptre et al., 2015). Recent range-wide population assessments have relied on extrapolation of occupancy models across different landscapes, with data centered on protected forests (Plumptre et al., 2021, 2016). This approach may miss localized effects, especially in areas not well represented by the data in the model such as at the edge of a species’ range, where environmental, anthropogenic, ecological, and behavioral patterns may diverge from the range center (Sexton et al., 2009).

Because great apes are rare, elusive, and inhabit dense forests, surveying them requires a proxy, typically the nests that adult individuals construct in new locations most nights (Kühl et al., 2008). To estimate absolute animal density, surveys commonly rely on “standing crop” nest counts along distance transects (Kühl et al., 2008). Field teams walk along randomly placed transect lines and record the perpendicular distance between each nest site and the transect line (Buckland et al., 2015). Given known nest production and decay rates, nest density can be converted to individual animal density (Buckland et al., 2015). However, nest decay rates vary significantly among habitats, seasons, height, and the plant materials used for construction, rendering density estimates based on rates calculated at another site or in a different habitat unreliable (Morgan et al., 2016; Romani et al., 2023; Tutin et al., 1995; Walsh and White, 2005). Survey-specific nest decay rates can be obtained with additional effort by locating and revisiting 50-150 fresh nests, ideally per species, habitat, and nest type stratum (Bessone et al., 2021; Kouakou et al., 2009; Laing et al., 2003). Similarly, the marked nest count method circumvents the issue of decay rates entirely, by re-walking transects and only considering fresh nests, but this not only requires additional time and effort but the appearance along transects of at least 60 fresh nests over the survey period (Kühl et al., 2008). Both of these options can be difficult, and even impossible, at low great ape density. Because the production and decay rates of other signs such as footprints, feeding remains, and dung are unknown, distance analyses are typically restricted to nest count data, leaving aside data on other signs even though such data are often collected together with nest data (*e.g.*, Plumptre et al., 2021).

A related challenge of the transect method in tropical forests is efficiency. Walking along a perfectly straight line and measuring precise perpendicular distances from that line is difficult and time consuming when the vegetation is dense. Field teams can often only walk a maximum of 1-2 km/day along line transects in central Africa (Caillaud, personal communication). When animal density is low, the sampling effort required to reach the recommended minimum of 60-80 groups of nests can be considerable (Kühl et al., 2008). Taking the example of Grauer’s gorillas, the exceptional 689 km, multi-year transect survey conducted by Plumptre et al (2021) only found 118 groups of nests. This means that observing 60-80 groups of nests required 350-467 km of transects. To increase sample size, authors sometimes analyze individual nests instead of groups (Carvalho et al., 2013; Haurez et al., 2014; Kouakou et al., 2009; Stokes et al.; 2010; van der Hoek et al., 2024). However, since nests from a given nest group are not statistically independent units, this approach underestimates the variance of nest density and the width of confidence intervals (Buckland et al., 2001; Haurez et al., 2014; Kouakou et al., 2009; Morgan et al., 2006; Stokes et al.; 2010). This leads to inflated type 1 error when animal densities are compared across sites, species, or time periods (Buckland et al., 2001).

Here, we propose an efficient and rapid method to survey wildlife distribution patterns that provides relative rather than absolute abundance estimates and is designed to be implemented by members of local communities. The resulting animal distribution maps and models of ecological covariates are not subject to errors induced by inaccurate nest decay rates, production rates, average group size estimates, or low sample size of groups of nests. Because this method, termed the “fast-transect” method, does not attempt to estimate absolute animal density, it can incorporate a variety of signs of animal presence in addition to nests to map animal distribution patterns, like dungs, feeding remains, tracks, and trails. Compared to transect analyses based on groups of nests, it uses more data, which increases the power of statistical analyses (Buckland et al., 2001). Importantly, fast-transect data collection does not require the same level of training as stranding crop nest counts, and can be performed by members of communities with limited access to formal education.

The 899 km^2^ Tayna Nature Reserve (TNR; **Fig. 1**) lies at the northeastern edge of the historical range of Grauer’s gorillas, at the border of the Lubero and Walikale territories in North Kivu, eastern DRC. The only published wildlife survey of TNR gorillas covered the central and southern portion of the reserve in 2003 (Mehlman, 2008). Two surveys were attempted since then, in 2006 and 2013, but neither was completed due to security issues (Nixon, 2013). Nevertheless, Plumptre et al. (2015) suggested a crash of the gorilla population in TNR by 89% between 2007 and 2013 based on nest encounter rates. Additional surveys are urgently needed to determine if a viable population of Grauer’s gorillas persists in TNR. If so, it would be one of the furthest north and east populations of Grauer’s gorillas left (aside from the few individuals at Mount Tshiaberimu, Virunga National Park, DRC; Iyer et al., 2023), which would warrant improved protection of the reserve itself, as well as the community conservation forests to the west, the Usala Corridor, that extends towards Maiko National Park 78 km away.

**Fig. 1.**
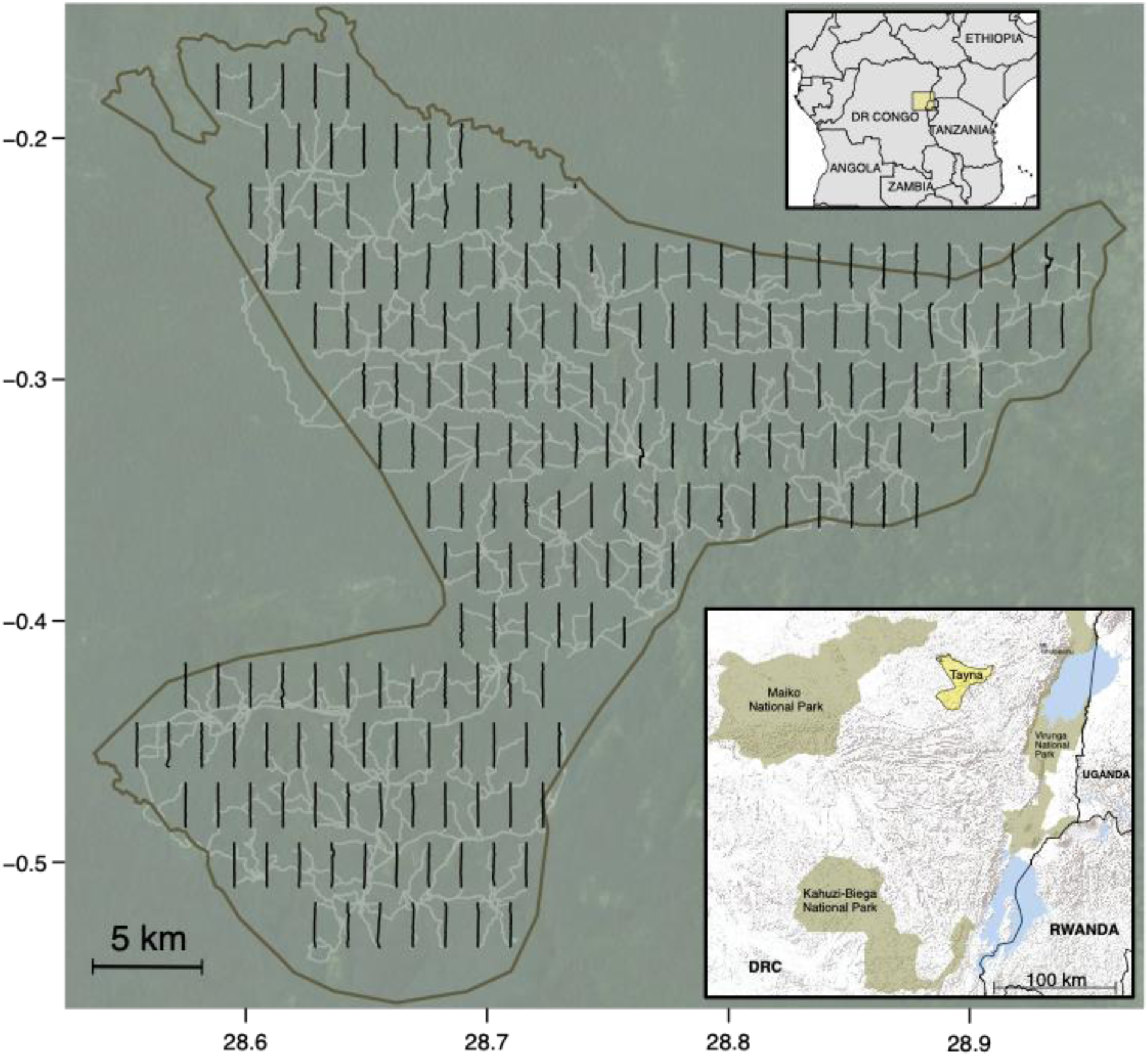
Situated in the eastern DRC, Tayna Nature Reserve (yellow, lower inset) lies in an ecological transition zone between the Albertine rift mountains to the east and the Congo Basin rainforest to the west. Black vertical lines correspond to the 196 line transects included in this survey, extracted from GPS tracks that recorded GPS points every 50 m. Grey lines correspond to paths walked to reach transects (grey lines). The background forest cover map is a composite satellite image from Google Earth (Maxar Technologies, Landsat/Copernicus 2020).

Here, we investigate the spatial distribution, terrain covariates, and impacts of anthropogenic disturbances on Grauer’s gorillas, eastern chimpanzees, guenons (*Cercopithecus spp.*), duikers (*Cephalophus spp.* and *Philantomba monticola*), and red river hogs (*Potamochoerus porcus*) in TNR. We conducted a 400-km fast-transect survey covering the entire reserve and leveraged generalized additive models to examine how wildlife distribution related to terrain and human activities. Data were collected by five teams of four people recruited from local communities living in and around TNR. We hypothesized that, if still present, Grauer’s gorillas and eastern chimpanzees would occur at lower densities near human settlements and forest activities such as artisanal mining and snare-trapping. Great apes are hunted for bushmeat (Ziegler et al., 2016), and even small-scale mining increases hunting pressure (Edwards et al., 2014; Spira et al., 2019). While it was unclear if large-scale trends applied locally, previous landscape-scale surveys found negative relationships between great ape density and distance to villages and mines (Plumptre et al., 2021, 2016). We expected the density of other wildlife species to also be impacted by human activities, but we predicted that the more resilient, faster-reproducing red river hogs and duikers would be less affected than slow-reproducing great apes and guenons (Estrada et al., 2017). Finally, we hypothesized that habitat suitability would vary across the heterogeneous terrain of TNR due to long-term human impacts and ecological gradients, such as shifts in plant community composition with altitude (Imani et al., 2016). As such, we predicted that wildlife distribution would relate to terrain, with different patterns for different species.

## 2. Methods

### 2.1. Study area

The Tayna Nature Reserve (TNR; 0° 20’ 36“ S, 28° 43’ 0” E; **Fig. 1**) was created in 2006 at the border of the Lubero and Walikale territories, North Kivu, eastern Democratic Republic of Congo. It encompasses 899 km^2^ of closed- and open-canopy primary forest. About 7% of its surface is agricultural land (Global Forest Watch, 2025). It lies in the transition zone between the Congo Basin and the Albertine Rift, generating a heterogeneous mix of high-altitude terrain up to 2,028 m above sea level (asl) stretching along the east of the reserve, with rugged hills and valleys near the center, and lower, flatter areas to the northwest dipping to 919 m asl (**Fig. 2A-B**). The forests of TNR extend continuously to the west to Maiko National Park. Vegetation is typical of low-to medium-altitude primary and secondary equatorial forest, though botanical surveys have not yet been undertaken. Rainfall averages 1,800-2,300 mm per year, with the rainiest season from October to November and a second rainy season from March to April (Congo Basin Forest Partnership, 2006). Temperature ranges from 13-34°C, with a daytime average of 19°C (Gorilla Rehabilitation and Conservation Education Center, unpublished data). The largest human settlements around TNR are to the northeast. Additional villages are found along the eastern and mid-western boundaries, and there are small dispersed settlements and encampments within the reserve (**Fig. 2C**). The survey was completed between September 12 and December 24, 2020.

**Fig. 2.**
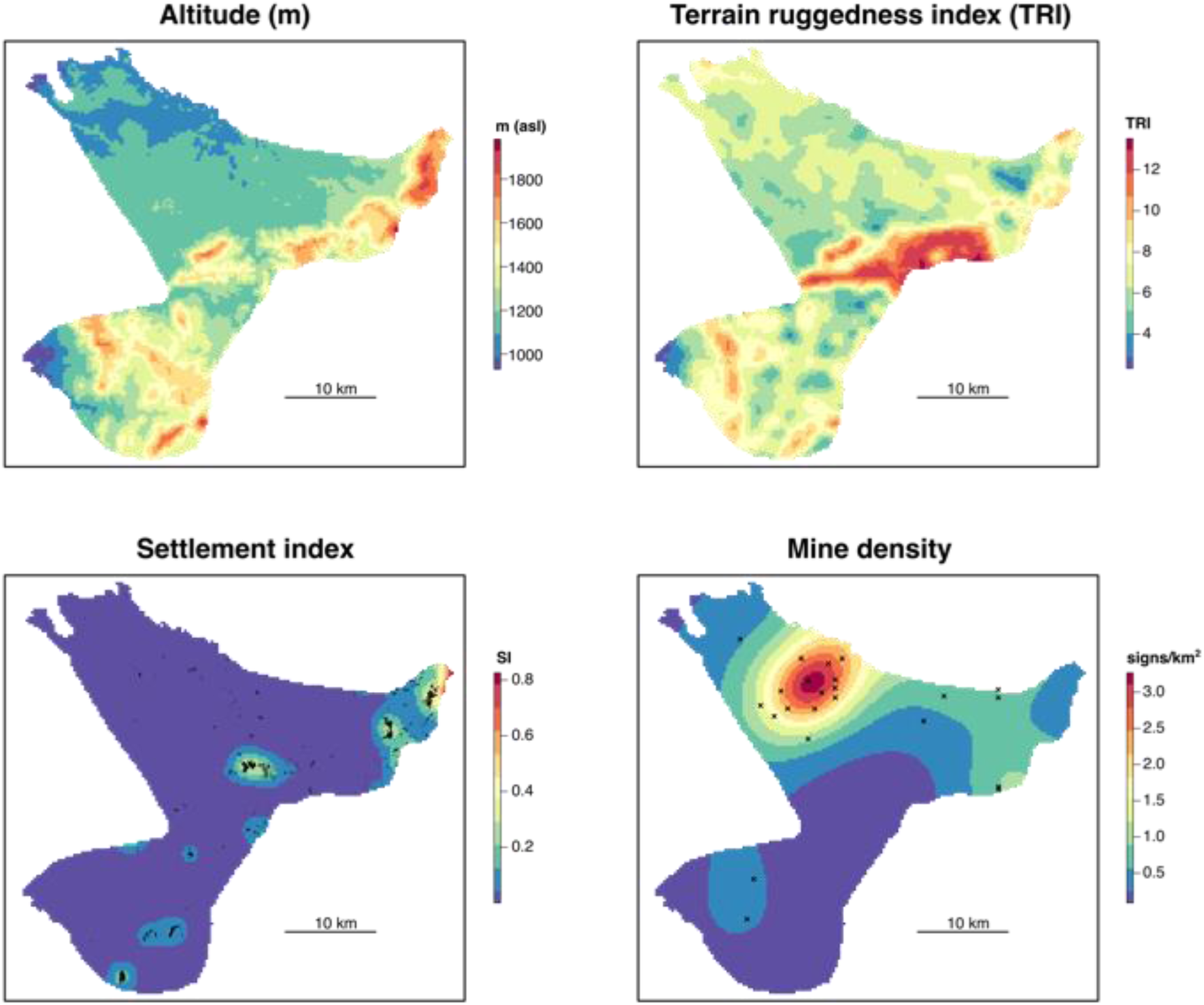
Maps of predictor variables for wildlife species distribution models in Tayna Nature Reserve. Altitude and TRI were calculated from a 30 m resolution digital elevation model (USGS, 2015; see Methods). Calculation of the Settlement Index and Mine density are described in the Methods section.

### 2.2. Fast-Transect Survey

Compared to distance transects (Buckland et al., 2015), fast-transects do not require measuring perpendicular distances between transect line and signs. Instead, observers are positioned on the transect line and on either side of it, sweeping a ∼100m strip of forest and searching for all signs of the species of interest. An important requirement for fast-transects is that the survey line be unbiased by habitat type (unlike reconnaissance surveys), so field teams follow a predefined compass bearing to the transect endpoint. The teams do not need to strictly adhere to this line, though, and can walk around localized obstacles such as patches of dense vegetation or fallen trees, as long as these deviations (typically less than 20m) are limited. This makes the data collection faster than traditional distance transects.

We designated 200 linear transects in a horizontally-staggered gridded pattern across the reserve (**Fig. 1**). Each transect was 2 km long, oriented north to south. Transects were spaced 1.5 km apart east to west and 750 m north to south. Each team comprised four members, all hired and trained locally: one team leader, one trail opener who walked ahead and cut minimally through the dense vegetation, and two observers on either side of the leader. The team leader carried a Garmin 64 handheld GPS, which automatically recorded points every 50 m. Waypoints for the start and end of each transect and any obstacles encountered were recorded in the GPS. The two observers “swept” about 50 m to the left and right of the leader for signs, covering a ∼100 m wide strip over each 2-km transect. As our goal was to estimate relative sign abundances, we did not need these two observers to sweep exactly 50 m of forest. Rather, they were trained to be consistent in the way they were walking and looking for signs. Observers remained within sight of the team leader, who recorded GPS points at sign locations. All signs of animal presence, including nest groups, footprints, feeding remains, dungs, burrows, vocalizations, and direct observations, were recorded and attributed to a particular species. We followed the method used by Plumptre et al. (2021) to assign great ape nest sites to gorillas or chimpanzees based on nest height, odor, dung, hairs, and trails. In addition, we recorded signs of human activities, such as camps, paths, snares, artisanal mines (operational and abandoned), logging, and crop fields. When signs of equivalent age, type, and species occurred within 50 m, only the first sign was considered, to reduce statistical non-independence issues caused by the clustering of signs produced by the same group of animals. Factors like rain could alter the decay time and detectability of signs, potentially inflating the between-transect variation in relative sign density. Our statistical approach explicitly modelled between-transect variation to limit the impact these factors might have had on the precision of our relative density estimates (Wood, 2017).

All data were collected strictly non-invasively in accordance with Congolese laws and regulations for wildlife research by local community members hired based on their tracking skills. Disturbance to wildlife during field work was negligible. Team members received extensive training in the fast-transect methodology at GRACE (Gorilla Rehabilitation and Conservation Education Centre), a research and conservation center located 7 km east of TNR.

### 2.3. Animal population distribution models

#### 2.3.1. Data preparation

To evaluate spatial heterogeneity in animal density in TNR, we modeled the probability of occurrence of animal signs using presence and absence points along transects and absence points generated randomly, as recommended by Barbet-Massin et al. (2012). To reflect the true sampling area , GPS track points were interpolated to a polyline and clipped to the start and end of each transect using QGIS. We excluded detours around obstacles and GPS errors (deviations over 50 m). Transect polylines were buffered by 50 m on each side, resulting in 100 m wide polygons covering the area “swept” by field teams along each transect. We retained only observations located within these polygons for our models, corresponding to a total survey area of 41.7 km^2^. We created *pseudo-*absence points by randomly placing 10,000 points within the polygons, as recommended for analysis of species distribution using generalized linear and additive models (Barbet-Massin et al., 2012). These *pseudo-*absence points were generated using function *spsample* in the ‘sp’ R package (Bivand et al., 2013; Pebesma and Bivand, 2005), corresponding to a density of 240 points per square kilometer of transect strip.

#### 2.3.2. Modeling framework

We opted for generalized additive models (GAMs) to model our presence/absence data because of their ability to incorporate nonlinear effects and two-dimensional (spatial) effects (Wood, 2017).

Models were fit in the R package ‘mgcv’ with a binomial distribution and logit link function. We used restricted maximum-likelihood to find the optimum smoothing parameters (Wood, 2018). Models were fit for each of the following animals: Grauer’s gorillas, eastern chimpanzees, guenons, duikers, and red river hogs. We modelled animal sign presence/absence in relation to four smoothed predictor variables: altitude, ruggedness, proximity to human settlements, and mining density (**Fig. 2**). All smooth terms were thin-plate regression splines with a basis dimension restricted to five to avoid overfitting. Because we had a large number of absence points (*n =* 10,000), model estimated probabilities (*p*) were very low (less than 0.01). At low *p* and high *n*, the binomial distribution converges towards the Poisson distribution (Poisson, 1837). As a consequence, our model’s probability estimates scaled perfectly with sign densities, and therefore, were proportional to animal densities. To add interpretability, we converted model-predicted probability of occurrence to relative density, measured in observed signs/km^2^, by multiplying the model prediction at each 300 x 300 m raster cell by the average sign density (total number of observed signs over 41.7 km^2^ of transect polygons), divided by the average predicted probability of sign occurrence across TNR. Hereafter we report modelling results in relative density (predicted number of observed signs/km^2^).

#### 2.3.3. Predictor Variables

To measure the altitude and terrain ruggedness in the region, we downloaded the Shuttle Radar Topography Mission 1 Arc-Second Global 30 m resolution digital elevation model (DEM) for the tile at 1°S, 28°E (USGS, 2015). Altitude (meters asl) was taken at the coordinates of each data point and pseudo-absence point. The terrain ruggedness index was computed as the mean of the absolute differences between the altitude at each DEM raster cell, in meters, and its eight adjacent cells (Wilson et al., 2007), averaged over a radius of one kilometer surrounding each data point. We chose one kilometer to reflect the average daily distance travelled by Grauer’s gorillas at low-altitude during the rainy season (van der Hoek et al., 2021a).

To evaluate the impact of human settlements, we calculated a “proximity index” that considers the distance to settlements as well as their relative size, since settlements in TNR ranged from camps to small villages. To do this, we drew polygons around settlements, fields, and camps visible in satellite imagery (Google Earth V7.3.3.7786, 2021) in TNR and the surrounding five km around it using QGIS (v.3.20). We converted the multi-polygon vector layer to a five-meter resolution raster using the R package ‘fasterize’ (R Core Team, 2023; Ross, 2020). Next, we converted the raster to a spatial points object, resulting in a grid of points every five meters over the settlement areas, using R functions *rasterToPoints* and *SpatialPoints* in the packages ‘raster’ package (Hijmans et al., 2013) and ‘sp’ (Pebesma and Bivand, 2005), respectively. We then interpolated the density of settlement points across TNR at the same resolution as the DEM (30 m) using kernel density estimation (KDE) with the R function *kernelUD* from package ‘adehabitatHR’, setting the smoothing parameter, *h*, to 1000 m (Calenge, 2006). Finally, we rescaled this settlement proximity function by dividing by its maximum value so it ranged between zero (far from settlements) and one (proximate to large settlements). This index captured the relative size of settlements as well as distance to them. Although the smoothing parameter (*h*) had a strong effect on KDE (Chen, 2017), its exact value did not have a major importance because the predictor variables in the GAM were allowed to have non-linear effects. Our only requirement for the KDE was that it captured some spatial variation in settlement density and gave more weight to larger settlements (Wood, 2017).

Along fast-transects, we observed signs of artisanal gold mining, consisting of one to a few small holes dug in the forest with an adjacent rudimentary camp that were either active or abandoned. To interpolate mining density across TNR, we fit a binomial GAM to the mining points and 10,000 pseudo-absence points using bivariate thin-plate splines to model the isotropic effects of latitude and longitude. This model predicted significant spatial variation in mining. The smoothed latitude-longitude predictor variable significantly influenced the probability of mining presence (*p* = 0.02). We converted these values to relative mining density by dividing them by their mean, and then multiplying resulting values by the observed mining density in transect strips (0.55 observed mining signs/km^2^) (**Fig. 2D**). We attempted the same approach to model the density of snares found along transects. However, latitude and longitude did not significantly influence snare density (*p* = 0.8). As a consequence, we could not include snare density as a predictor variable in our animal distribution models (Bryn et al., 2021).

#### 2.3.4. Testing and accounting for spatial autocorrelation

Spatial autocorrelation is important to consider in species distribution modelling (Gaspard et al., 2019). If left unaccounted for, it violates model assumptions of nonindependence and can cause spurious association with covariates (Dormann et al., 2007). Our data collection protocol only considered successive observations over 50 m apart, which likely reduced the risk of spatial autocorrelation. To determine whether this was sufficient, we evaluated spatial autocorrelation using semi-variograms of GAM residuals for points within five km of one another, with the R package ‘gstat’ (Pebesma, 2004). We did not detect significant spatial autocorrelation for any of the animal species considered (**Fig. S1**). Thus, we did not incorporate spatial autocorrelation in GAMs.

#### 2.3.5. Model diagnostics

We used two strategies to assess the fit of GAMs. First, we mapped predicted relative density for each species using the *predict* function in the ‘raster’ package, taking as predictor variables a raster stack of the terrain and anthropogenic variables in our model (Hijmans et al., 2013). We plotted observations of animal signs on top of this map to visually inspect model fit. Second, we assessed model fit by binning the predicted probabilities from the GAM into 25 equal-sized groups. Bins had equal counts to avoid differences in variance, so their widths varied. To evaluate alignment between predictions and observations, we calculated the observed presence rate for each bin (i.e., the proportion of points with detected animal sign) and standard error, and plotted these against the mean predicted probability.

## 3. Results

### 3.1. Fast-transect survey

The five survey teams completed 196 of the 200 planned transects. Four transects were inaccessible. Teams covered one or two transects per day and seven to twelve transects per two-week patrol over 22 total patrols, equating to a survey effort of 162 work days, excluding travel and rest days (76 actual days because of contemporaneous deployment). The average pace of fast-transects was 2.4 km/day (SD = 0.08 km/day across teams).

We recorded 87 signs of Grauer’s gorilla and 151 signs of eastern chimpanzees both on and off transects in TNR. Signs included nest sites, tracks, trails, feeding remains, dung, vocalizations, and visual (for chimpanzees) observations. Among these signs, 41 gorilla signs and 119 chimpanzee signs were found on fast-transects (**Table 1**). There were 0.99 gorilla signs observed per km^2^ of transect strip and 2.87 chimpanzee signs/km^2^, or 0.10 gorilla signs per km of fast-transect walked and 0.30 chimpanzee signs/km. In addition to great apes, we recorded signs of cercopithecine monkeys, duikers, at least two pangolin species (*Smutsia gigantea* and *Phataginus tricuspis* and/or *P. tetradactyla*), red-river hog, brush-tailed porcupine (*Atherurus africanus*), forest buffalo (*Syncerus caffer nanus*), aardvark (*Orycteropus afer*), and leopard (*Panthera pardus*) (**Table 1**). There were no signs of forest elephants (*Loxodonta cyclotis*).

**Table 1.** Species identified along transects in Tayna Nature Reserve (TNR). Animal signs were recorded on 196 transects that covered 41.7 km^2^ of the 899 km^2^ of TNR.

| Wildlife species | Seen | Heard | Tracks, trails, dung,<br>and food remains | Sleeping sites,<br>*i.e.*, nest sites or<br>burrows, etc |
| --- | --- | --- | --- | --- |
| *Gorilla beringei graueri* | 0 | 1 | 24 | 16 |
| *Pan troglodytes schweinfurthii* | 1 | 18 | 45 | 55 |
| *Cercopithecus spp.* | 65 | 108 | 28 | - |
| *Cephalophus spp.* &<br>*Philantomba monticola* | 11 | 0 | 380 | 18 |
| *Potamochoerus porcus* | 0 | 0 | 223 | 5 |
| *Atherurus africanus* | 1 | 1 | 196 | 7 |
| *Phataginus spp.* &<br>*Smutsia gigantea* | 0 | 0 | 12 | 34 |
| *Syncerus caffer nanus* | 0 | 0 | 19 | - |
| *Panthera pardus* | 0 | 0 | 5 | - |
| *Orycteropus afer* | 0 | 0 | 1 | 5 |

During the survey in TNR, both on and off transects, we found 292 gorilla nests in 40 nest sites (including 93 nests in 16 sites along transects) and 280 chimpanzee nests in 69 nest sites (including 209 nests in 55 sites along transects; **Fig. S2**). Excluding 23 solitary nests (15 attributed to chimpanzees and 8 to gorillas), the average nest group size was 8.89 (SD = 5.72) for gorillas and 4.91 (SD = 2.88) for chimpanzees (**Fig. S3**).

### 3.2. Population distribution models

#### 3.2.1. Grauer’s Gorillas

The relative density of gorilla signs observed in TNR was positively associated with terrain ruggedness (GAM, *edf* = 2.88, χ^2^ *=* 14.50, *p =* 0.003), but not related to altitude (*edf* = 1.00, χ^2^ *=* 1.84, *p =* 0.18). It was negatively associated with the anthropogenic factors settlement index (*edf* = 1.00, χ^2^ *=* 9.98, *p =* 0.002) and mine density (*edf* = 1.00, χ^2^ *=* 12.37, *p* < 0.001; **Table 2**). Model fitted values aligned visually with off-transect observations of gorilla signs (**Fig. 3a**). There was a clear positive relationship between observations and model predictions, indicating that the model was able to capture population distribution (**Fig. S4a**).

**Table 2.** Generalized additive models of great ape sign density in TNR

| ***Gorilla beringei graueri*** |  |  |  |  |
| --- | --- | --- | --- | --- |
| **Term** | ***Edf*** | **Ref df** | **$\chi^2$** | ***p*** |
| Thin-plate smoothed terms |  |  |  |  |
| Altitude | 1.000 | 1.001 | 1.838 | 0.175 |
| Terrain ruggedness index | 2.879 | 3.285 | 14.504 | **0.003** |
| Settlement index | 1.000 | 1.000 | 9.977 | **0.002** |
| Mining density | 1.000 | 1.000 | 12.373 | **<0.001** |
| *n* = 10041 |  |  |  |  |
| *Deviance explained* = 19.6% |  |  |  |  |
| ***Pan troglodytes schweinfurthii*** |  |  |  |  |
| **Term** | ***Edf*** | **Ref df** | **$\chi^2$** | ***p*** |
| Thin-plate smoothed terms |  |  |  |  |
| Altitude | 1.000 | 1.001 | 2.886 | 0.089 |
| Terrain ruggedness index | 1.918 | 2.371 | 45.327 | **<0.001** |
| Settlement index | 1.000 | 1.000 | 4.124 | **0.042** |
| Mining density | 1.883 | 2.298 | 5.936 | **0.047** |
| *n* = 10119 |  |  |  |  |
| *Deviance explained* = 8.63% |  |  |  |  |

**Fig. 3.**
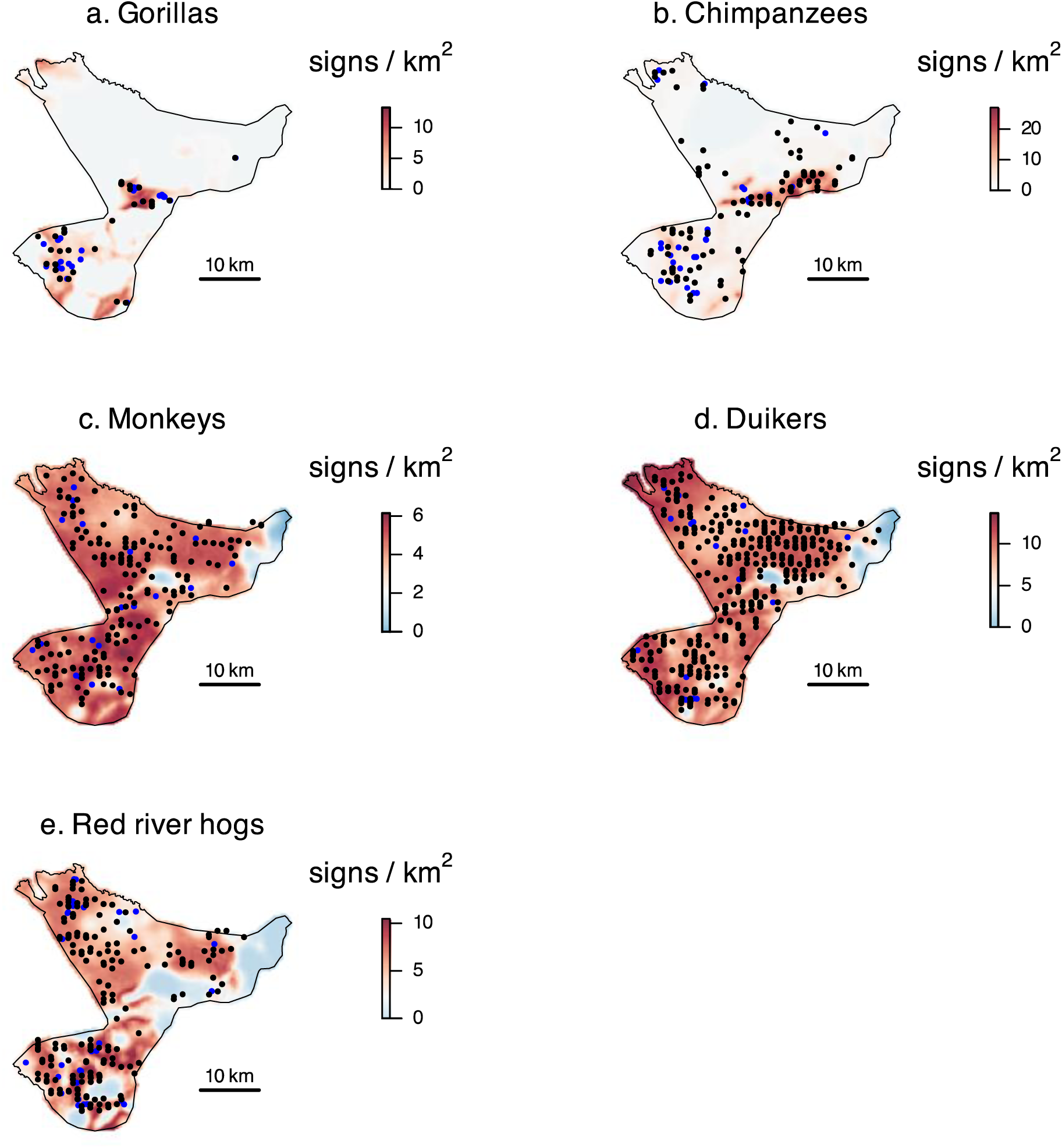
Predicted relative densities of animal populations in observed signs/km^2^ in TNR based on GAM probability of occurrences (see Methods). The average density of signs of gorillas, chimpanzees, monkeys, duikers, and red-river hogs on transects were 0.99, 2.87, 4.15, 8.52, and 4.46 per km^2^, respectively. Animal sign observations on transects are marked in black circles, which were the presence points in the model. Off-transect sign observations are shown in blue.

The model showed that gorilla sign relative density was maximized at ruggedness values above 7.5, which was above the average ruggedness of TNR (7.04, SD = 1.71, range = 3.04 – 12.90, median = 6.60), and plateaued thereafter (**Fig. 4a**). The settlement index was negatively associated with gorilla sign density, with gorilla sign density declining to below average at values just above the median settlement index in TNR (0.006, median = 0.004; **Fig. 4a**). The effect of mine density was negative, and no signs of gorillas were found in mining hotspots (**Figs. 2 and 3**).

**Fig. 4.**
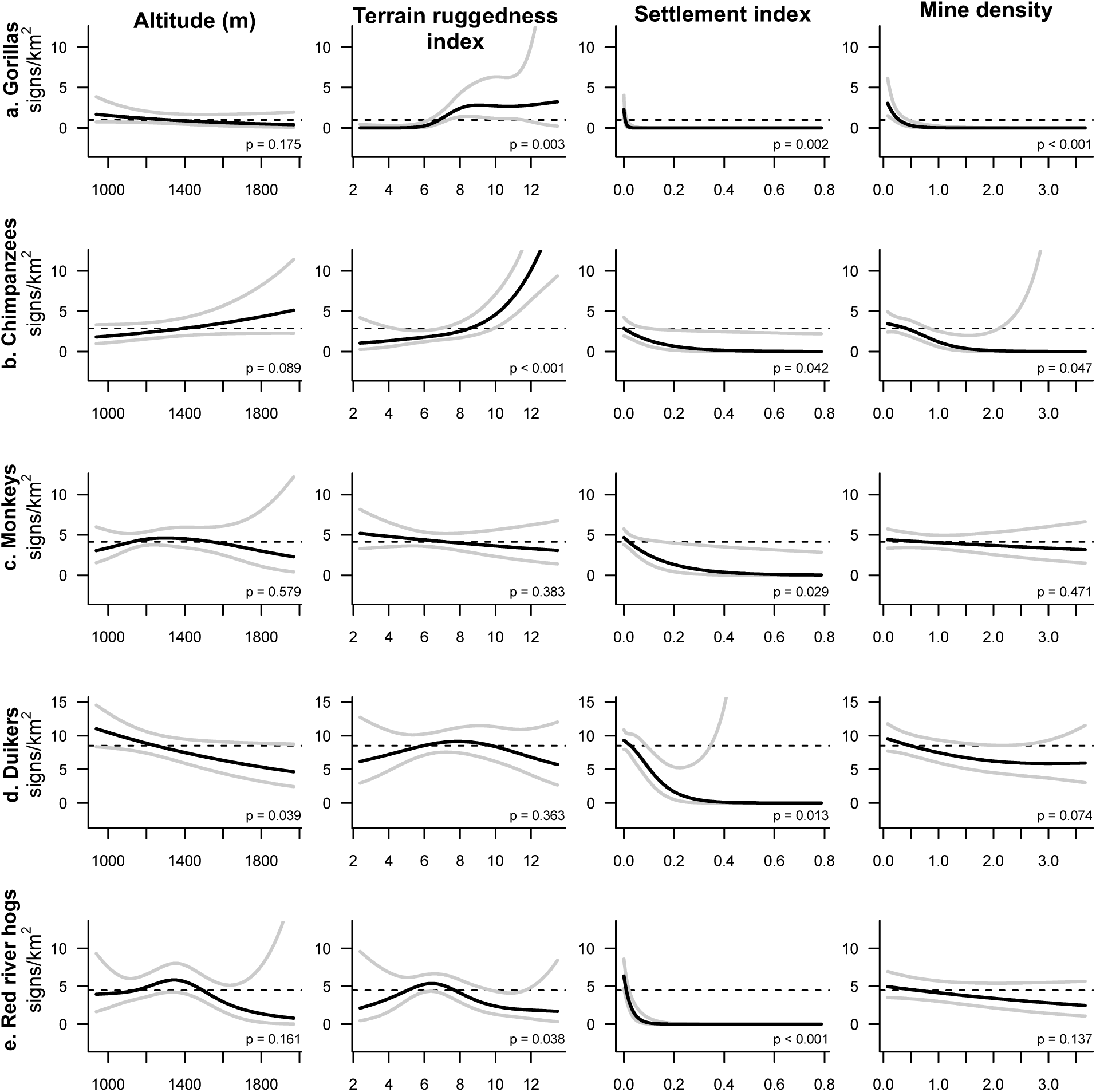
Partial effects of altitude, terrain ruggedness, settlement proximity, and mine density on predicted animal sign density in observed signs per km^2^ (see Methods). The dashed line shows the mean sign density for each population. Gray lines show the upper and lower 95% confidence intervals around the main effect prediction (black line).

#### 3.2.2. Eastern chimpanzees

The density of chimpanzee signs was positively associated with terrain ruggedness (GAM, *edf* = 1.92, χ^2^ *=* 45.33, *p <* 0.001) but not altitude (*edf* = 1.00, χ^2^ *=* 2.89, *p =* 0.09), and negatively associated with the settlement index (*edf* = 1.00, χ^2^ *=* 4.12, *p =* 0.04) and mining density (*edf* = 1.88, χ^2^ *=* 5.94, *p =* 0.05; **Table 2**; **Fig. 3b**). Chimpanzee sign density was predicted to peak at maximal ruggedness (TRI > 12; **Fig. 4b**). Compared to gorilla signs, chimpanzee signs were more uniformly distributed across TNR, including in the far north where great ape signs were never attributed to gorillas (**Fig. 4a**). The fitted GAM for observed chimpanzee signs predicted well the distribution of observations (**Fig. S4b**).

#### 3.2.3. Monkeys

The survey recorded sightings, vocalizations, prints, and food remains from guenons. From direct observations only, we detected at least five different species: red-tailed monkey (*Cercopithecus ascanius*), blue monkey (*C. mitis*), owl-faced monkey (*C. hamlyni*), mountain monkey (*C. lhoesti*), and Dent’s mona monkey (*C. denti*). Considering signs of all monkey species combined, they were less spatially variable than those of great apes (**Fig. 3c**) and were not statistically associated with variation in altitude (GAM *edf* = 1.88, *χ^2^ =* 1.66, *p* = 0.58), ruggedness (*edf* = 1.00, *χ^2^ =* 0.76, *p* = 0.38), nor mine density (*edf* = 1.00, *χ^2^ =* 0.52, *p* = 0.47). Settlement proximity had a significant negative effect on the relative density of monkey signs observed (*edf* = 1.003, *χ^2^ =* 4.77, *p* = 0.03; **Fig. 4c**). As expected from the relatively even sign distribution, the predictive capability of the GAM for monkey signs was low (**Fig. S4c**).

#### 3.2.4. Duikers

Signs and sightings of commonly-called “red duikers” (duikers of similar size with similar footprints and dung, such as black-fronted duiker, *Cephalophus nigrifrons*, Weyn’s duiker, *C. weynsi*, bay duiker, *C. dorsalis*, and white-bellied duiker, *C. leucogaster*), yellow-backed duiker (*Cephalophus silvicultor*), and blue duiker (*Philantomba monticola*) were observed widely across TNR (**Fig. 3d**).

Duiker signs consisted of prints, trails, food remains, and dung. The significant factors influencing the distribution of duiker species’ signs combined were settlement index (GAM *edf =* 1.77, *χ^2^ =* 9.27, p = 0.01) and altitude (*edf =* 1.00, *χ^2^ =* 4.28, p = 0.04). More duiker signs were recorded at low values of both. Relative density appeared to decline at high mining density, but this effect was not significant (*edf* = 1.56, *χ^2^ =* 4.53, p = 0.07; **Fig. 4d**). There was a clear positive relationship between observations and model predictions, indicating that the model was able to capture population distribution (**Fig. S4d**).

#### 3.2.5. Red-river hogs

Red-river hogs left signs including trails, prints, dung, and food remains on transects. They were heard and seen once (three individuals) off-transect. The settlement index had a strong negative effect on the distribution of observed red river hog signs (GAM *edf =* 1.000, *χ^2^ =* 20.58, p < 0.001; **Figs. 3e and 4e**). Terrain also mattered, with sign density peaking just below the average ruggedness (*edf =* 2.64, *χ^2^ =* 8.95, p = 0.04; **Fig. 4e**). There was a clear positive relationship between observations and model predictions, indicating sufficient predictive power of the model (**Fig. S4e**).

## 4. Discussion

We used a novel field data collection method, termed “fast-transects”, to survey Tayna Nature Reserve (TNR) in north Kivu, DRC, in 2020. Our survey identified populations of Critically Endangered Grauer’s gorillas and Endangered eastern chimpanzees. Observed signs of gorillas and chimpanzees were positively associated with increasing terrain ruggedness, but not altitude, and negatively influenced by proximity to human settlements. Gorilla signs also showed a negative association with artisanal mining in TNR. Signs of duikers, monkeys, and red river hogs were distributed more homogeneously throughout the reserve than either of the great apes. Nevertheless, all species groups were negatively impacted by proximity to human settlements. The survey also detected signs from other wildlife species, including forest buffalo, leopard, pangolins, and aardvark.

### 4.1. Comparison with prior surveys

This is the first survey to cover the entirety of TNR. We found that the Grauer’s gorilla population in TNR is restricted to the rugged mountains and valleys in the center and south of the reserve. Gorillas were present in this sector in the early 2000s, based on partially-completed surveys (Mehlman, 2008; Nixon, 2013; Plumptre et al., 2015). The gorilla population seems to have persisted since then, at least in the central and southern sector. This may indicate that poaching of gorillas has been minimal since the legal establishment of TNR in 2006, although firm conclusions cannot be drawn in the absence of data from the 2000s from the vast northern sector of the reserve (Mehlman, 2008; Nixon, 2013; Plumptre et al., 2015). Like gorilla signs, chimpanzee signs were predominantly found in the center and south of TNR, but, contrastingly, chimpanzee signs were also found in the northern sector. Neither the distribution nor abundance of chimpanzees was reported in previous surveys (Mehlman, 2008; Nixon, 2013), but the encounter rate with nests appeared to have declined by over 80% from 2006 to the uncompleted 2013 survey (Nixon, 2013; Plumptre et al., 2015). Other species had not been formally surveyed previously, though the 2013 survey team in the center of TNR reported that no monkeys were seen or even heard (Nixon, 2013).

### 4.2. Influence on terrain on species distribution

Gorilla and chimpanzee signs in TNR were positively associated with ruggedness, but not altitude. Rugged terrain may provide foraging opportunities or refuge against predators and competitors, or the preference may reflect inaccessibility to humans. This finding may be a general characteristic of Grauer’s gorillas and eastern chimpanzee in this region, as Plumptre et al. (2021) found a positive association between terrain steepness, a variable that likely correlates with terrain ruggedness, and great ape abundance in Kahuzi-Biega National Park and the Oku Community Reserve. In the tropics, plant diversity tends to be maximized in mountainous ridges and valleys (Baldeck et al., 2013). Great apes have diverse diets, comprising terrestrial herbaceous vegetation, seasonal fruit, roots, and more (*e.g.,* Basabose, 2002; Michel et al., 2022; van der Hoek et al., 2021b), and they may select areas with diverse plant communities where fallback foods and preferred foods co-occur within short distances. An alternative possibility is that great apes were extirpated from flatter areas due to hunting, or that they avoid such areas because of current or past human activities.

We did not detect an effect of terrain ruggedness or altitude on guenon relative density. If the negative association between flat terrain and great ape signs reflects hunting pressure, guenons may rebound quicker due to their larger population sizes and faster reproductive rates (Ingram et al., 2021), or their presence in risky habitats may reflect territorial exclusion or source-sink dynamics, where “source” rugged habitats are already at ecological carrying capacity (Novaro et al., 2000). It is also possible that terrain variability does not influence food resources for Cercopithecine monkeys or their perception of safety as they are more arboreal than great apes (Clark and Kaplin, 2024; Tuyisingize et al., 2023).

Importantly, in our analysis we combined monkey species because of identification uncertainty for indirect signs, potentially smoothing over interspecific variation (Chapman et al., 2004). Duiker and red river hog signs were more common in low-altitude and less rugged areas, respectively. Little is known of the ecological preferences of either animal group in the region, so more research would be needed to formulate and test hypotheses explaining this pattern (but see Bobo et al., 2025; van der Hoek et al., 2023).

### 4.3. Influence of human presence & activities on species distribution

Human settlements and signs of artisanal mining negatively affected the relative density of Grauer’s gorillas and, to a slightly lower extent, eastern chimpanzees. Negative associations between great apes and both small-scale mining and villages have been reported in the Grauer’s gorilla range at large (Plumptre et al., 2021), as well as for closely related great apes in west central Africa (Morgan et al., 2018; Ordaz-Németh et al., 2021; Strindberg et al., 2018). Notably, chimpanzees were found in the northern sector of TNR in areas where gorillas were not found. This may reflect their flexible multi-male fission-fusion social system and smaller group sizes than gorillas, which could render them more tolerant to hunting pressure. Other aspects of eastern chimpanzee behavior demonstrate their adaptability. For example, chimpanzees living adjacent to croplands have been observed raiding crop fields at night (Krief et al., 2014).

Relative densities of monkeys, duikers, and red river hogs were also lower near human settlements. Regionally, duikers are the most commonly consumed bushmeat, with red river hogs second (Batumike et al., 2021; Mbete et al., 2011; van Vliet et al., 2015). Artisanal mining did not appear to have a significant effect on these taxa. This could reflect lower hunting pressure near mines than near more permanent settlements.

Local community culture, including preferences, dietary taboos, traditional forest products, and hunting tactics, result in prioritized offtake of different wildlife species and could drive the observed pattern (Malimbo et al., 2020). While cultural transitions are likely eroding this source of variation (Chausson et al., 2019; Mbete et al., 2011), traditional hunting practices are still important in this region.

The long-term absence of great apes from a significant part of the TNR forest may have consequences on ecosystem functions (Abernethy et al., 2013). For example, in Nigeria, primate-dispersed tree seedlings are rarer in areas where hunting is not regulated, compared to protected areas (Effiom et al., 2013). Whether the structure of the TNR forest is affected by the heterogeneous distribution of apes requires further research.

### 4.4. Methodological limitations

Fast-transects provided an efficient, rapid means to map the distribution of wildlife activity at the population level. When the density of the type of animal sign used for a distance analysis is low, making derived population densities too imprecise, fast-transects provide a practical solution. Relative to distance transects, fast-transects enable more data to be collected in less time, reducing financial and logistical costs to conservation organizations and reducing the risk that insecurity or unforeseen circumstances prevent survey completion. If we had used the traditional method of standing crop nest counts on distance transects, we would have only been able to analyze data from 16 gorilla nest sites and 55 chimpanzee nest sites, at best. As Kühl et al (2008) recommend a minimum of 60-80 nest sites to calculate great ape density using this method, it is clear that relying on nests only would have prevented us from testing most of our hypotheses regarding great ape distribution. Instead, the fast-transect method allowed us to analyze a total of 160 great ape signs. Combined with powerful non-linear models, the larger dataset allowed us to detect significant effects of terrain and human activity on great ape distribution.

The main limitation of the fast-transect method is its inability to estimate absolute animal densities and population sizes (Buckland et al., 2015). However, even distance transects based on standing crop nest counts often fail at providing reliable absolute estimates, due to their reliance on published nest decay rates from other geographical areas. A variation of the distance transect method requiring marking observed nests and revisiting transects regularly reduced this bias, but this requires a substantial data collection effort often incompatible with large geographical areas and impossible when nest density is too low (Kouakou et al., 2009; Spehar et al., 2010). Because we did not attempt to estimate absolute densities, we could analyze all sign types. Since each sign has particular biases (*e.g.,* dung; Kamgaing et al., 2018; and vocalizations; Croes et al., 2007), combining multiple sign types helps build a more accurate model of the distribution of wildlife rather than a particular behavior (*e.g.*, nesting or alarm-calling).

### 4.5 Conservation perspectives & conclusions

The persistence of gorillas and chimpanzees in TNR underscores the importance of community land protection in the eastern Albertine Rift biodiversity hotspot. Our survey results provide a much-needed baseline for future surveys in TNR. Repeated fast-transects, currently underway, will track changes in animal relative density patterns, letting managers cater development, sensitization, and education programs to specific communities and identified threats. TNR now has field teams collecting data on the movement patterns and diet of gorilla groups in the central rugged region, where this survey identified multiple groups (Mbeke, personal communication). These continuous activities offer long-term employment to community members living in closest proximity to wildlife (Mbeke, personal communication).

As a peripheral population, TNR gorillas are more exposed to the risks of habitat fragmentation and genetic isolation (van der Valk et al., 2018). Maintaining connectivity between TNR and other wildlife populations is critical, beginning with the forested landscape that stretches westward toward Maiko National Park. The status of great apes and other wildlife populations within this remote area, the Usala Corridor, remains uncertain. Building on this study, a fast-transect survey is currently underway in this region. Usala features heterogeneous terrain, yet it has a different pattern of anthropogenic pressures compared to TNR. A comparative study could help disentangle whether the association between great apes and rugged terrain observed in TNR is due to ecological factors, human avoidance, or historical extirpation.

Globally, forest fragmentation is increasing at an alarming rate, causing biodiversity loss and disrupting ecological processes (Taubert et al., 2018). The present survey is only the first step towards the long-term protection of TNR. Moving forward, the continuation of monitoring activities and the development of research will maintain the engagement of local communities and support long-term conservation outcomes.

## Acknowledgments

We are grateful to the many field assistants and trackers hired from the local villages surrounding Tayna Nature Reserve, without whom this study would not have been possible, including the following field team leaders: Arnold Kakule Vahwere, Affable Kakule Pilipili, Faustin Kambale Kinaba, Celestin Kasereka Kunemutumba, Moise Kalingini, Faustin Muhindo Kibwana, and Evariste Muhindo Katsongo. We thank the GRACE DRC team for managing ongoing survey logistics, in particular Augustin Kambere Mbangi. We thank Finley for his support. Two anonymous reviewers provided valuable suggestions. This work was funded by the Disney Conservation Fund’s Reverse the Decline Initiative, the Margot Marsh Biodiversity Foundation, and the University of California, Davis.

## CRediT authorship contribution statement

**Alice Michel:** Data curation, Formal analysis, Investigation, Methodology, Software, Visualization, Writing – original draft, Writing – review & editing. **Jackson Kabuyaya Mbeke:** Conceptualization, Funding acquisition, Methodology, Project administration, Resources, Supervision, Writing – review & editing. **Benezeth Kambale Visando:** Data curation, Investigation, Project administration, Resources, Supervision, Writing – review & editing. **Sonya Kahlenberg:** Conceptualization, Funding acquisition, Methodology, Project administration, Supervision, Writing – review & editing. **Katie Fawcett:** Funding acquisition, Project administration, Supervision, Writing – review & editing. **Damien Caillaud:** Conceptualization, Data curation, Formal analysis, Investigation, Methodology, Project administration, Supervision, Validation, Writing – original draft, Writing – review & editing.

## Data Availability

All data and code are available in the A.P.E.S. database (Kühl et al., 2007).

## Appendix A: Supplementary Materials

Separate Word document containing supplementary figures.

