## Supplemental Information for "Terrain ruggedness and human activities influence the distribution of Grauer’s gorillas (*Gorilla beringei graueri*) and eastern chimpanzees (*Pan troglodytes schweinfurthii*) in the Tayna Nature Reserve, Democratic Republic of Congo"

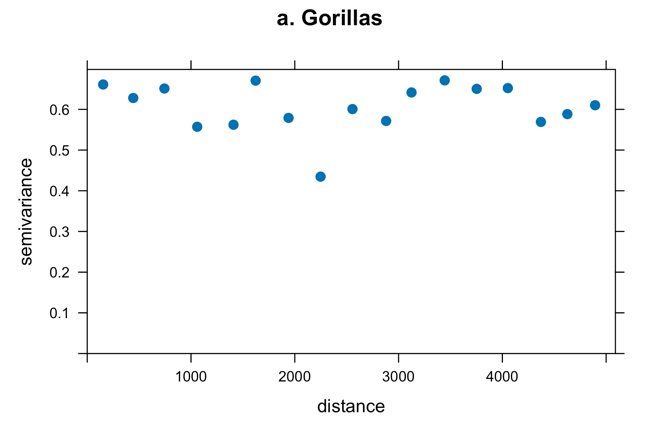

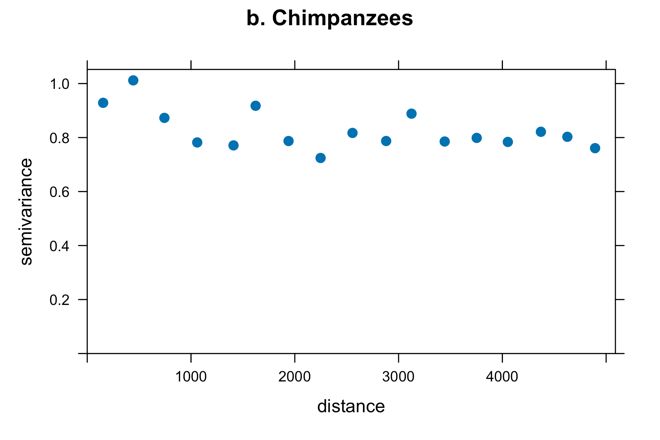

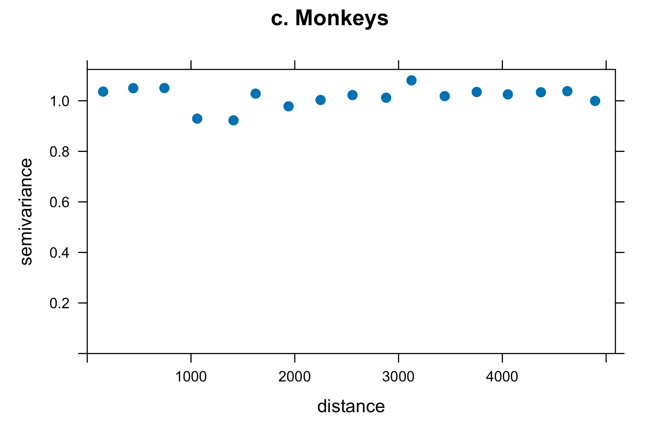

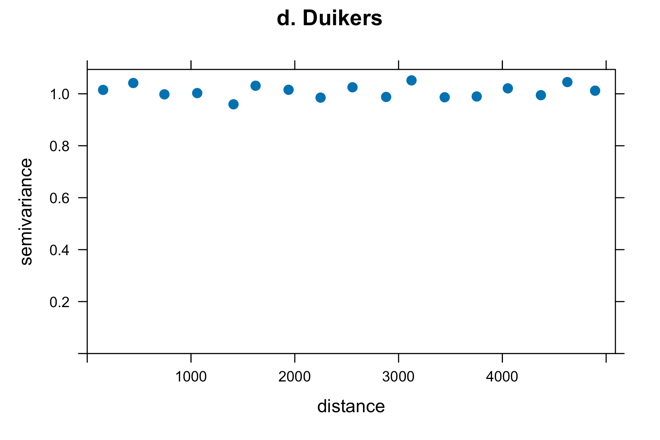

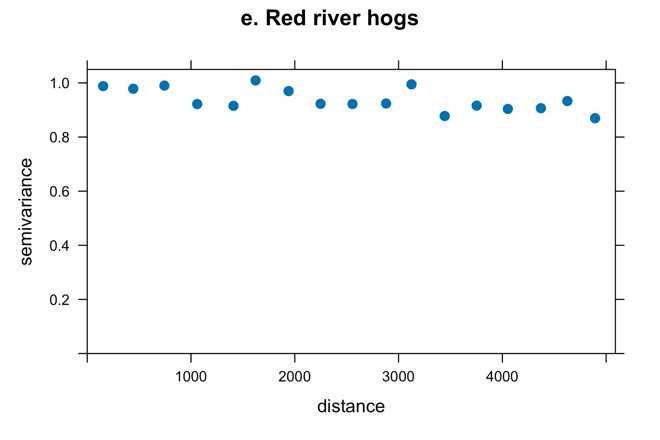

**Supplemental Figure A1.** Semivariograms of GAM model residuals. Spatial autocorrelation was not detectable for any species modeled.

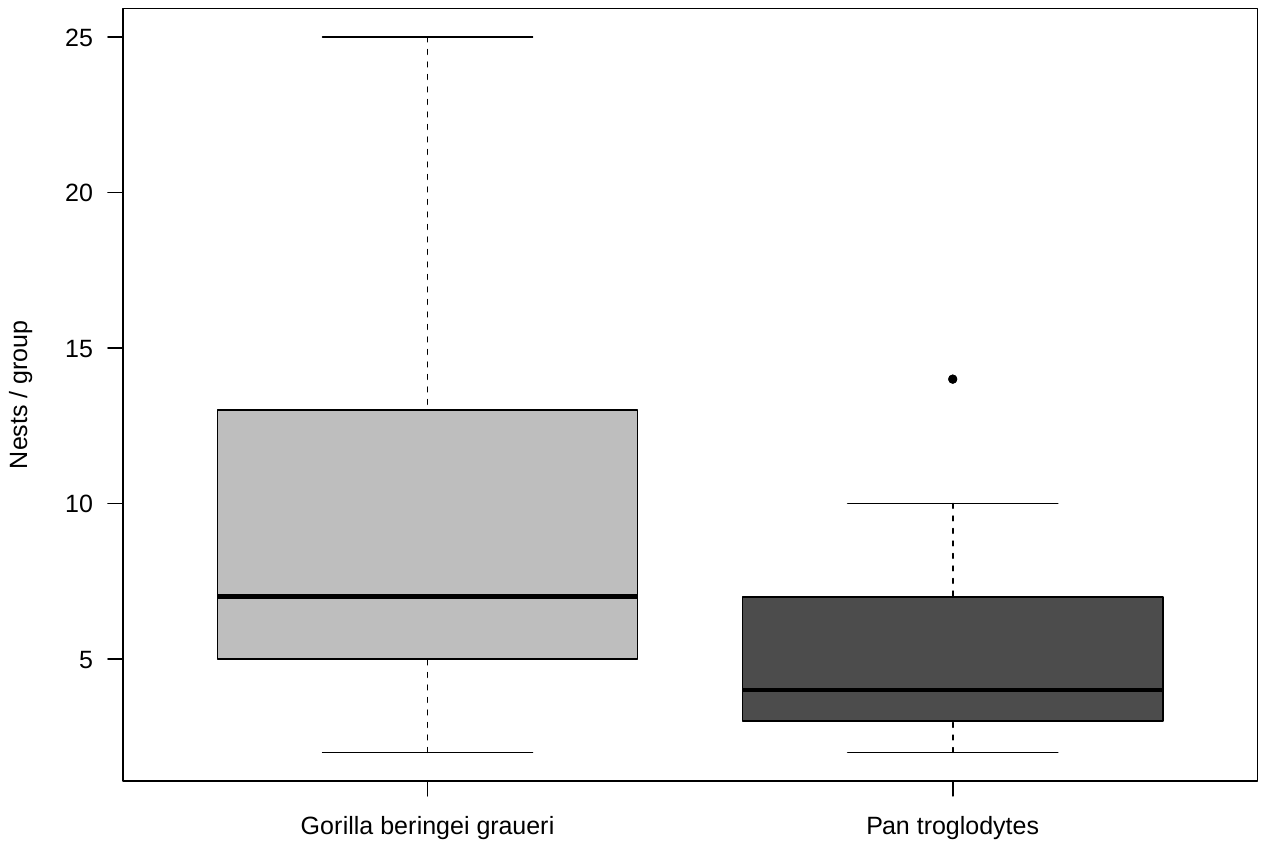

**Supplemental Figure A2.** Excluding solitary nest sites, great ape nest sites attributed to gorillas (n=32) had more nests than those attributed to chimpanzees (n = 54; mean 8.88 vs. 4.91; GLM estimate = -0.23, se = 0.07, z = -3.20, *p =* 0.001).

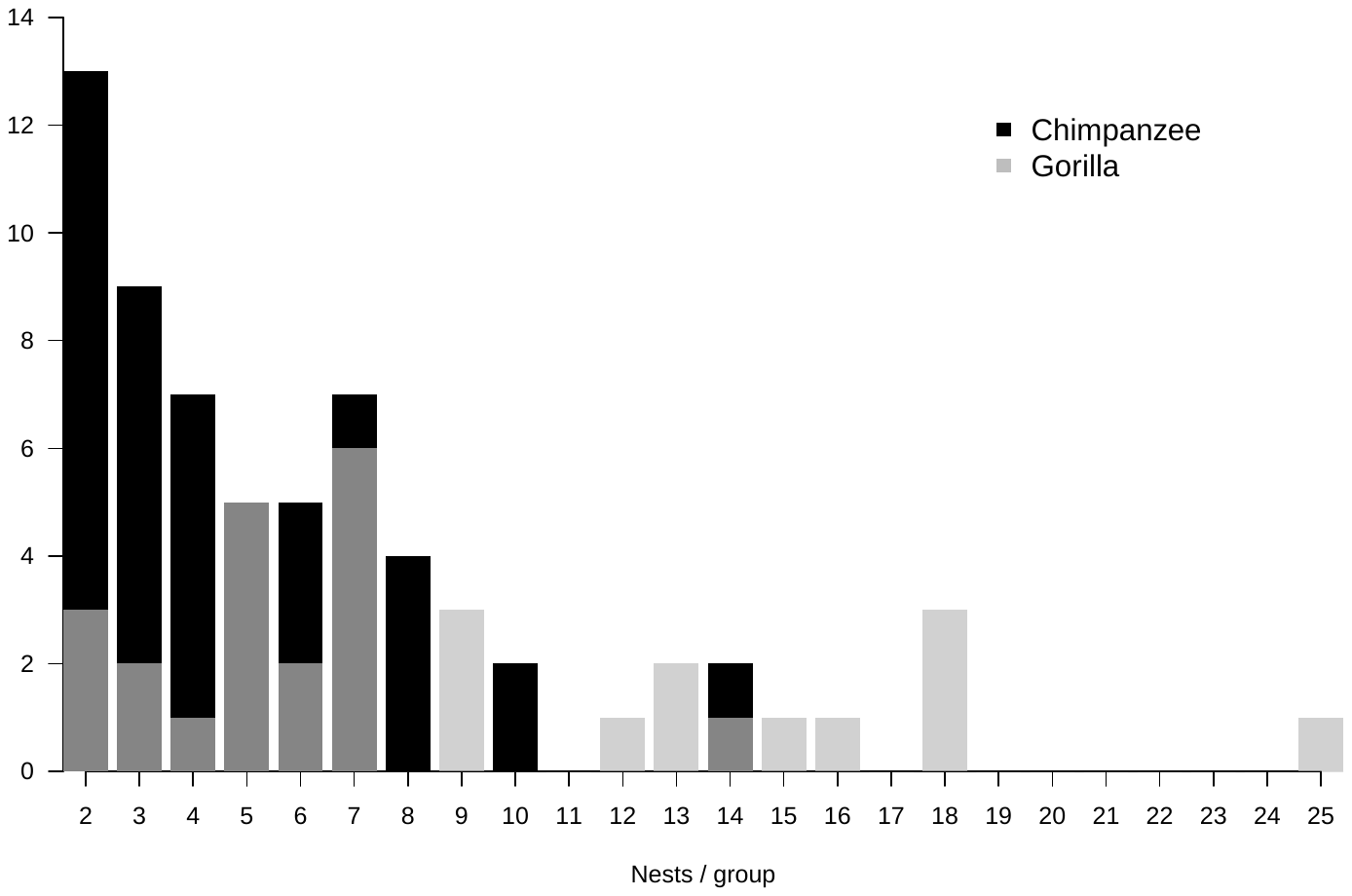

**Supplemental Figure A3.** The distribution of the number of nests per nest site attributed to chimpanzees and gorillas. Solitary nest sites are excluded.

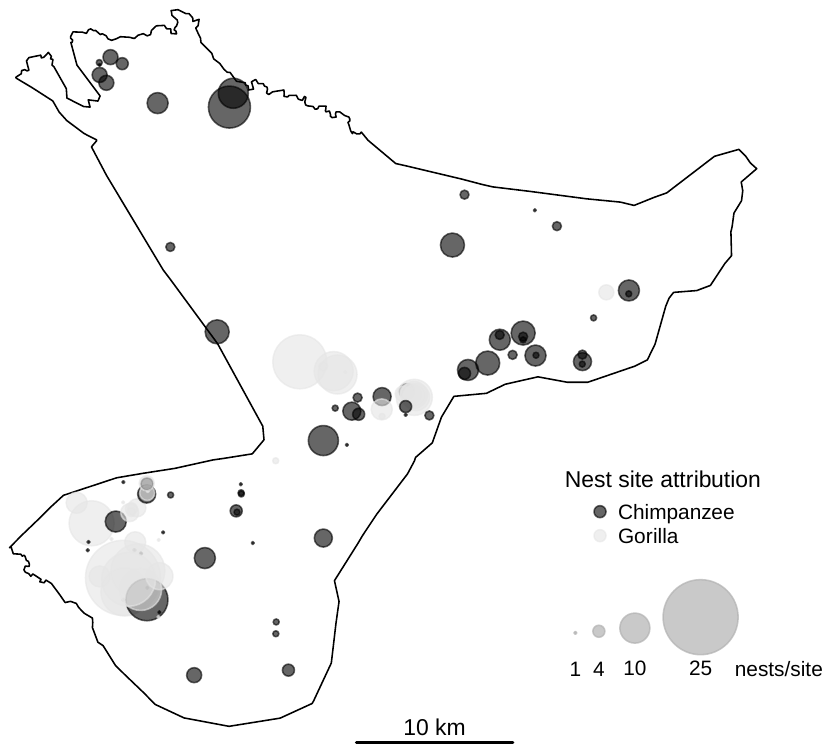

**B**

**A**

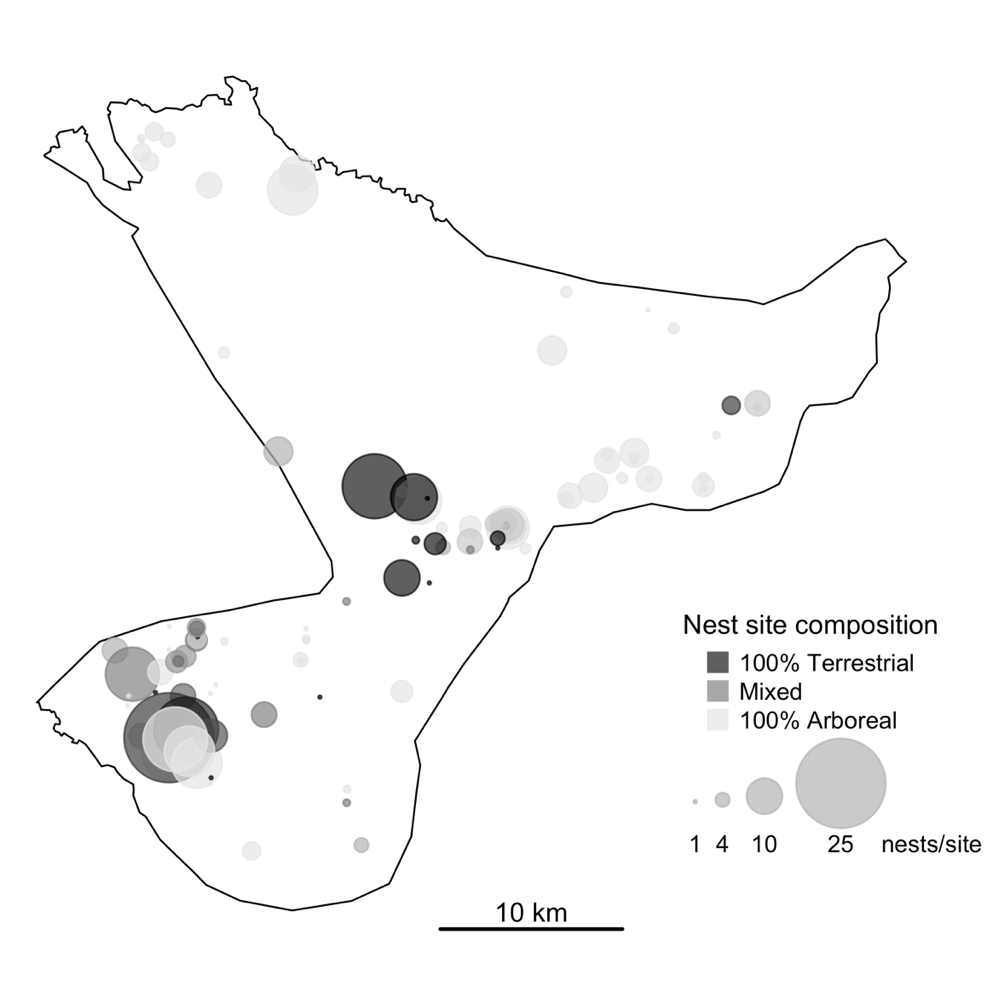

**Supplemental Figure A4.** Great ape nest sites on and off transect in TNR. The circle size is scaled by number of nests per site. (A) and (B) show the same data, but in (A), the color shows species, with dark grey nest sites attributed to chimpanzees and light grey to gorillas. In (B), the color scale shows the percent of the total nests in the trees per site, from all (white) to none (dark grey).

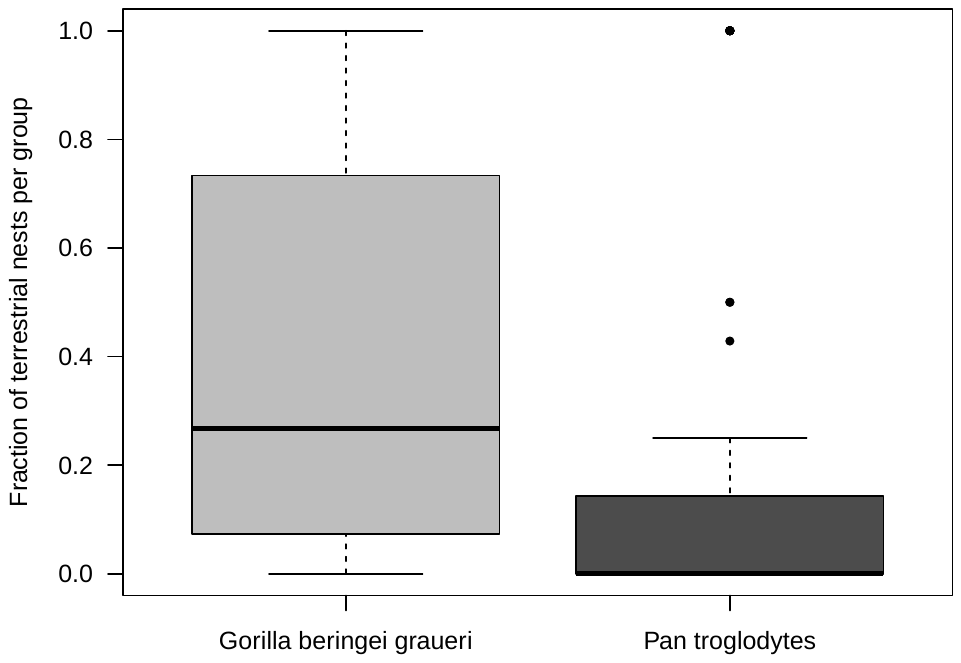

**Supplemental Figure A5.** Fraction of terrestrial nests out of total nests per site was higher for nest sites attribute to gorillas than chimpanzees (mean 39% vs. 18%; GLM estimate = -2.09, se = 0.83, z = -2.51, *p =* 0.01). In general, gorilla nest sites tended to vary more in terms of arboreality whereas chimpanzees tended to nest more in trees in TNR, with notable outliers.

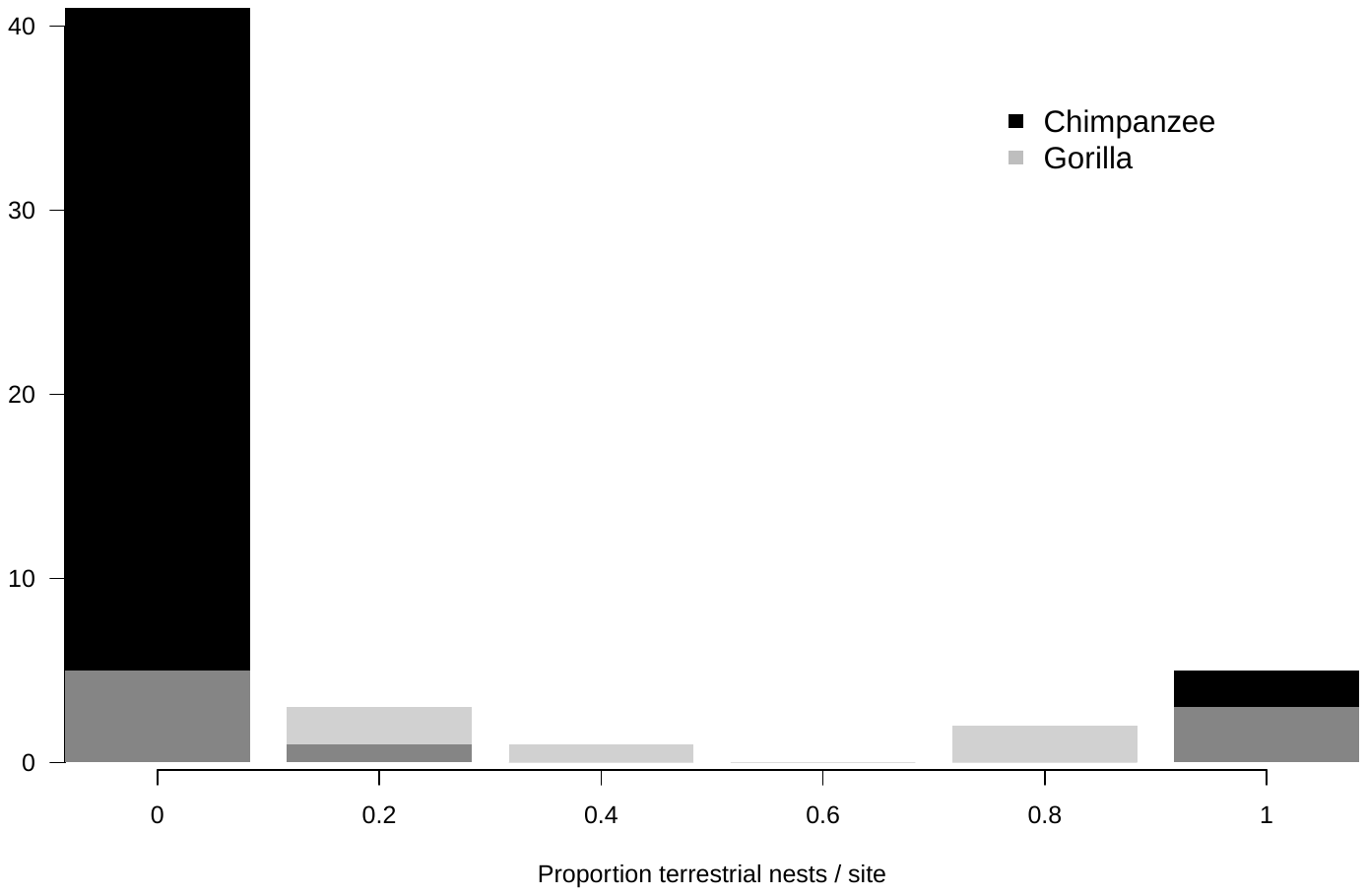

**Supplemental Figure A6.** The distribution of the fraction of ground, or terrestrial, nests out of total nests per site of the two different great ape species in TNR.

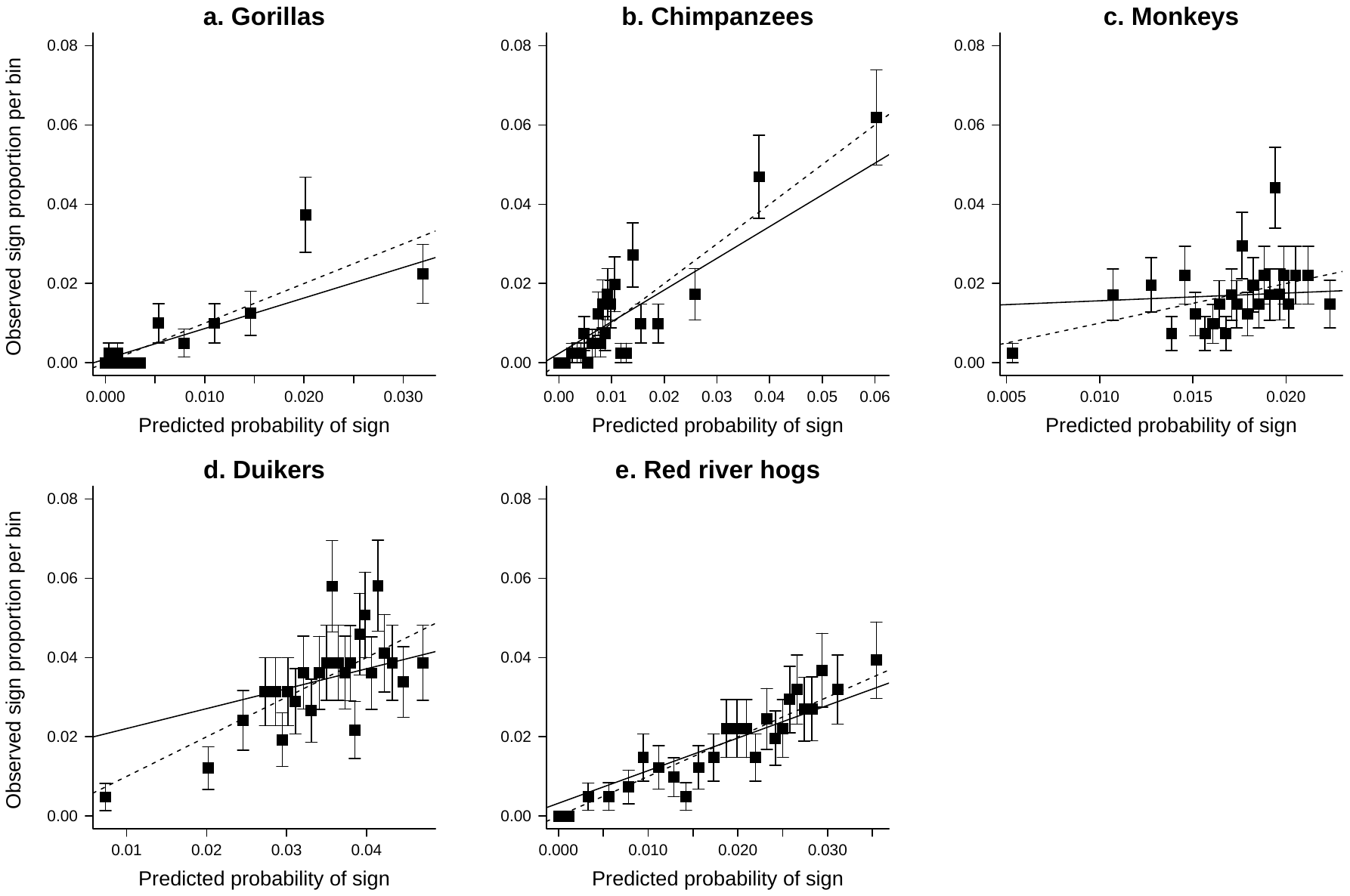

**Supplemental Figure A7**. Model predictive capacity shown by binned predictions for each GAM for each of (a) gorillas, (b) chimpanzees, (c) monkeys, (d) duikers, and (e) red river hogs. Fitted values from each GAM were placed into 25 quantile groups, taking the average value per bin as the predicted probability of sign shown on the x-axis. The observed sign proportion per bin shown on the y-axis is the fraction of presence to absence points in that quantile group. The dashed line indicates a one-to-one perfect fit, and the solid line is a linear regression of the actual fit of observed to predicted.
